# Tolerance and dependence to Δ^9^-tetrahydrocannabinol in rhesus monkeys: activity assessments

**DOI:** 10.1101/499152

**Authors:** David R. Schulze, Lance R. McMahon

**Author notes:** Correspondance: Lance R. McMahon, Ph.D., Department of Pharmacodynamics, College of Pharmacy, University of Florida, 1345 Center Drive, Building JHMHSC Room P1-20, P.O. Box 100487, Gainesville, Florida 32610.

## Abstract

*Cannabis* withdrawal upon discontinuation of long-term, heavy *Cannabis* use is reported in humans; however, methods to establish the nature and intensity of cannabinoid withdrawal, especially directly observable signs, have not been widely established. This study quantified activity in the home cage of rhesus monkeys, and examined the extent to which activity can be used to quantify tolerance to and dependence on Δ^9^-tetrahydrocannabinol (Δ^9^-THC). Home-cage activity was measured in one group that received Δ^9^-THC (1 mg/kg s.c.) every 12 h (i.e., chronic Δ^9^-THC), and a second group that received Δ^9^-THC (0.1 mg/kg i.v.) once every 3 days (i.e., intermittent Δ^9^-THC). Treatment was temporarily discontinued in the chronic Δ^9^-THC group and the effects of rimonabant and Δ^9^-THC were examined in both groups. Activity counts were highest during the day (lights on 0600–2000 h) and were lower at night. Rimonabant (0.1–3.2 mg/kg i.v.) dose-dependently increased activity (maximum 20-fold) in the chronic Δ^9^-THC group, but did not significantly alter activity in the intermittent Δ^9^-THC group. Δ^9^-THC (0.32–3.2 mg/kg i.v.) dose-dependently decreased activity counts (maximum 4-fold) in both groups, but was somewhat more potent in the intermittent as compared with the Δ^9^-THC group. Discontinuation of Δ^9^-THC treatment resulted in an immediate (i.e., within 24 h) and time-related increase in activity. Resumption of Δ^9^-THC treatment (1 mg/kg/12 h) produced hypoactivity that was no longer evident within 9 days of treatment. The time-related increase in home-cage activity upon abrupt discontinuation of chronic Δ^9^-THC treatment, as well as the effects of rimonabant to increase activity in monkeys receiving chronic, but not intermittent, Δ^9^-THC treatment, are consistent with signs of physical dependence on Δ^9^-THC in primates.

## INTRODUCTION

Long-term cannabis use results in dependence evidenced by a withdrawal syndrome after abrupt discontinuation of use (Jones 1983; Vandrey and Haney 2009 for reviews). Cannabis withdrawal consists primarily of self-reported symptoms including anger, anxiety, depressed mood, irritability, headaches, tension, stomach pain, and strange dreams. Directly observable signs of cannabis withdrawal are typically less prevalent than self-reported symptoms. Directly observable signs include aggression, sleep difficulty, decreased food intake, weight loss, and sweating. Cannabis withdrawal signs and symptoms typically emerge within 24 h after abrupt discontinuation of cannabis use and subside within 1–2 weeks. Cannabis withdrawal, although not life-threating, is clinically significant inasmuch as individuals use cannabis to avoid or alleviate withdrawal (Budney et al. 1999).

Nonhuman primates have been used in studies of dependence and withdrawal to a variety of drugs including Δ^9^-tetrahydrocannabinol (Δ^9^-THC), the drug primarily responsible for the behavioral effects of cannabis, although Δ^9^-THC dependence has not been unanimously reported. Observable withdrawal signs in rhesus monkeys were not evident after discontinuation of Δ^9^-THC treatment (2 mg/kg/day for 30 days; Harris et al. 1974). Cardiovascular function in rhesus monkeys did not show evidence of a withdrawal syndrome after discontinuation of Δ^9^-THC treatment (2 mg/kg/day for 3 weeks; Fredericks et al. 1981). Observable signs of withdrawal also did not emerge upon termination of daily treatment with another cannabinoid agonist (levonantradol) in rhesus monkeys (Young et al. 1981). However, in contrast to these negative findings, abrupt discontinuation of Δ^9^-THC (2 mg/kg/day for 3 weeks) increased some observable behaviors above baseline (Fredericks and Benowitz, 1980), and there was a time-related disruption in operant responding for food when continuous i.v. administration of Δ^9^-THC (0.05 mg/kg/h) was discontinued (Beardsley et al. 1986). Collectively, these studies not only illustrate the mixed results typical of pre-clinical studies examining Δ^9^-THC dependence, but they also underscore the relatively mild nature of cannabis withdrawal as compared to withdrawal from other drugs such as ethanol and opioids.

CB_1_ receptor-selective antagonists such as rimonabant have been useful in pre-clinical studies of cannabinoid dependence; however, the use of rimonabant has not been free of limitations. Rimonabant surmountably antagonized many of the effects of cannabinoid agonists (e.g. Compton et al., 1996). Moreover, rimonabant alone produced behavioral effects including hyperactivity, scratching, and wet-dog shakes in rodents (Compton et al., 1996; Aceto et al., 1996), and headshaking and increased heart rate in rhesus monkeys (Stewart and McMahon, 2010). Although rimonabant produced many of these effects under conditions of chronic Δ^9^-THC treatment, their presence in the absence of cannabinoid treatment suggests that such effects are unrelated to dependence and withdrawal. On the other hand, rimonabant produced some effects during chronic Δ^9^-THC treatment but not in the absence of Δ^9^-THC treatment, including headshakes in rodents (Aceto et al., 1996) and discriminative stimulus effects in rhesus monkeys (Stewart and McMahon, 2010). Moreover, some of the effects of rimonabant in Δ^9^-THC treated animals are observed upon abrupt discontinuation of Δ^9^-THC treatment (McMahon and Stewart, 2010), indicating that those effects are signs of Δ^9^-THC withdrawal.

One goal of the present study was to examine the extent to which movement (i.e., motor activity) of individually housed rhesus monkeys in the home cage is sensitive to Δ^9^-THC dependence and withdrawal. The Δ^9^-THC treatment (1 mg/kg/12 h s.c.) used here was the same as that used to establish and maintain a rimonabant discriminative stimulus (Stewart and McMahon, 2010). A second and related goal was to compare abrupt discontinuation of Δ^9^-THC treatment to administration of rimonabant during Δ^9^-THC treatment. Locomotor activity was assessed during Δ^9^-THC treatment, after treatment was abruptly discontinued, and for several weeks after resumption of Δ^9^-THC treatment, as well as after various doses of rimonabant during Δ^9^-THC treatment. Because Δ^9^-THC treatment had been ongoing for years at the time these studies were conducted, a separate group of monkeys was used to control for Δ^9^-THC treatment. Monkeys in the second group had discriminated a relatively small dose (0.1 mg/kg i.v.) of Δ^9^-THC administered once every 3 days on average for years. While the second group was not a negative control, the marked difference in Δ^9^-THC treatment dose and frequency was expected to provide a sufficient control for the impact of Δ^9^-THC treatment on the effects of rimonabant.

## MATERIALS AND METHODS

### Subjects

Five adult rhesus monkeys (*Macaca mulatta*; three female and two male) received 1 mg/kg/12 h of Δ^9^-THC s.c. Five additional adult rhesus monkeys (three male and two female) received 0.1 mg/kg of Δ^9^-THC i.v. on average once every 3 days. Monkeys were housed separately on a 14-h light/10-h dark schedule (lights on at 0600 h), were maintained at 95% free-feeding weight (range 5.6–10.1 kg) with a diet consisting of fresh fruit (apples, bananas, and oranges), peanuts, and primate chow (High Protein Monkey Diet, Harlan Teklad, Madison, WI). Monkeys received food at 1600 h each day and water was available continuously in the home cage. Monkeys received non-cannabinoids and cannabinoids in previous studies (Stewart and McMahon 2010; McMahon 2011). Experiments were conducted in accordance with the Institutional Animal Care and Use Committee, The University of Texas Health Science Center at San Antonio, and with the “Guidelines for the Care and Use of Mammals in Neuroscience and Behavioral Research” (National Research Council 2011).

### Surgery

A catheter (heparin coated polyurethane, od = 1.68 mm, id = 1.02 mm, Instech Solomon, Plymouth Meeting, PA) was inserted 5 cm into a femoral or subclavian vein under anesthesia with ketamine (10 mg/kg i.m.) and isoflurane (1.5–3.0% inhaled via facemask). Suture silk (coated vicryl, Ethicon Inc., Somerville, New Jersey) secured the catheter to the vessel and was used to ligate the section of the vessel adjacent to the catheter insertion. The opposite end of the catheter was attached to a vascular access port (Mida-cbas-c50, Instech Solomon), which was located s.c. in the mid-scapular region of the back.

### Apparatus

Activity was monitored in stainless-steel home cages measuring 33 inches wide × 27 inches deep × 32 inches high. The activity monitor was an ActiCal Activity Monitor 64 K Memory Waterproof (Philips Respironics Mini-Mitter Company, Inc., Bend OR). The monitor was attached to an Actiwatch Animal Case (Phillips Respironics), which was fastened to the Primate Products collar worn around the neck. The Actical Data Acquisition algorithm involves an electric sensor, which generates a voltage when it undergoes a change in acceleration. Thirty-two times per second, the filtered, amplified voltage is converted to a digital value, is used to adjust a running baseline value, and is added to a one second accumulated value. Every second, the one-second sum is divided by four and added to a resultant accumulated value for the epoch. At the end of each epoch, the accumulated activity value is compressed into an 8-bit value and stored in Actical memory. The data are downloaded by Windows software, the 8-bit values are decompressed to 15-bit raw activity counts, and the Actical-specific calibration constant is applied to the raw activity counts, resulting in calibrated activity data.

### Drugs

Rimonabant and Δ^9^-THC (100 mg/ml in absolute ethanol; The Research Technology Branch of the National Institute on Drug Abuse, Rockville, MD) were dissolved in a mixture of 1 part absolute ethanol, 1 part Emulphor-620 (Rhodia Inc., Cranbury, NJ), and 18 parts physiologic saline and were administered in a volume of 0.1 ml/kg for subcutaneous administration and 0.1–1 ml/kg for intravenous administration. For the largest dose (3.2 mg/kg) of rimonabant, the vehicle was 1 part absolute ethanol, 2 parts Emulphor-620, and 7 parts physiologic saline.

### Discontinuation study

Three monkeys were used for the discontinuation study. Monkeys receiving daily Δ^9^-THC (1 mg/kg/12 h s.c.) had been treated for at least two years before the discontinuation study. One week before discontinuation they were fitted with an activity monitor and data was collected to establish an activity baseline. During discontinuation of Δ^9^-THC treatment, monkeys received vehicle instead of Δ^9^-THC starting at 6:15 PM; the following day was defined as the first full day of discontinuation (Day 1). Monkeys received vehicle for the next 19 days (Days 2–20) and at 6:15 AM the following day (Day 21). Δ^9^-THC treatment was resumed at 6:15 PM and the following day was defined as the first full day of resumed Δ^9^-THC treatment. To administer 1 mg/kg of subcutaneous Δ^9^-THC or vehicle at 6:15 AM and 6:15 PM, monkeys were pulled toward the front of the home cage via a retractable back wall, a process requiring less than 1 min.

### Intravenous Δ^9^-THC and rimonabant

Two groups of monkeys (n=5 per group) were used. One group received daily Δ^9^-THC (1 mg/kg/12 h) and the other group received intermittent Δ^9^-THC (0.1 mg/kg i.v. every 3 days on average). For intravenous injections of Δ^9^-THC and rimonabant, monkeys were removed from the home cage and were seated in chairs (Model R001, Primate Products, Miami, FL) at 12:10 PM. A dose of drug or vehicle was administered and monkeys were returned to the home cage at 12:15 PM. Activity was measured until 2:15 PM.

### Treatment order

For both the discontinuation and the i.v. Δ^9^-THC/rimonabant studies, a given treatment was not studied in more than one monkey on any given day. That is, the Δ^9^-THC discontinuation experiment was initiated on different days for each monkey and only one monkey received a given dose of Δ^9^-ΤHC or rimonabant on any given day.

### Data analyses

An activity count was generated every 15 s and these were cumulated (i.e., 1, 2, or 24 h) for data presentation and analysis. For the discontinuation study, daily activity patterns for individual monkeys were plotted by cumulating activity in 1-h bins and showing these data across a 24-h period during daily Δ^9^-THC treatment, after discontinuation, and upon its resumption. These data were expressed as a mean of the 5 days during Δ^9^-THC treatment immediately preceding discontinuation, the first 5 days of discontinuation, the last 5 days of discontinuation, and the first 5 days of resumption of daily Δ^9^-THC treatment. For statistical analysis of data generated during the discontinuation experiment, cumulative activity over 24 h was expressed as a percentage of the individual control defined as the average of the 5 days of activity immediately preceding discontinuation of Δ^9^-THC treatment. A significant difference was defined as mean activity that fell outside the largest and smallest of the 95% confidence limits determined for any of the 5 days immediately preceding discontinuation.

To examine the effects of intravenous Δ^9^-THC (0.1–3.2 mg/kg) and rimonabant (0.1–3.2 mg/kg), data were cumulated for 2 h post-injection and were plotted and analyzed as a percentage of the control. For each monkey, activity counts were cumulated over 2 h following administration of vehicle; data from two separate vehicle tests were averaged for further analysis. Straight lines were simultaneously fit to individual dose-response data with linear regression (Prism version 5.0 for Windows, GraphPad Software Inc., San Diego, CA). The slopes and intercepts of the dose-response functions determined in the chronic and intermittent Δ^9^-THC groups were compared to determine whether a single line was sufficient to describe the data (i.e. the two dose-response functions were not significantly different) or two lines were needed to describe the data. Activity counts were averaged per individual, expressed as a percentage of control, and plotted as a mean ± S.E.M. for the group.

## RESULTS

### Daily Δ^9^-THC treatment: control activity counts

In monkeys that had received 1 mg/kg of Δ^9^-THC every 12 h for at least two years, activity counts were highest during the light period and lowest during the night (Figure 1; each panel shows data from an individual). Activity was variable among monkeys with two monkeys showing lower overall activity (Figure 1 top and middle) than a third monkey (Figure 1 bottom; note the different range on the ordinate). When expressed as cumulative activity counts per h, the highest activity occurred 3–5 pm. The individual maximum values were 1987, 3143, and 7977 counts per h (Figure 1 top, middle, and bottom respectively, circles). In contrast, the activity counts during the night were as low as 200, 181, and 106 in each respective monkey. In general, a period of inactivity began at 7 pm (2 h before dark onset) and ended at 5 am. Cumulative 24-h activity was relatively stable during the 5 days immediately preceding discontinuation of Δ^9^-THC treatment, i.e., daily activity did not deviate more than −2% (day 3) to +12% (day 4) of the 5-day running mean. For the 95% confidence limits calculated for each day of the 5-day running mean, the maximum upper limit was 168% (day 5) and the minimum lower limit was 54% (day 2).

**Figure 1.**
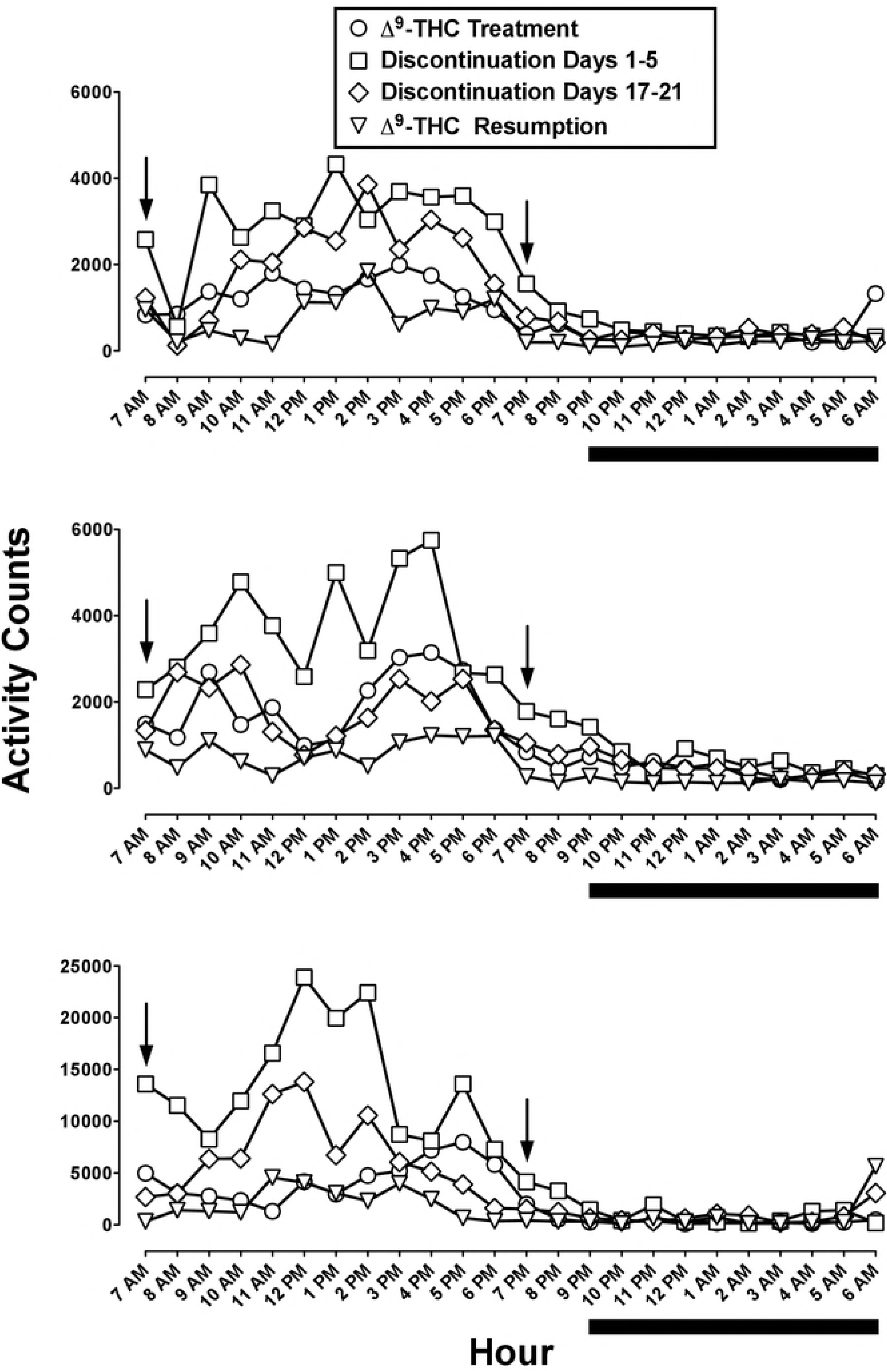
Absolute activity counts cumulated in 1-h bins and expressed for each h of the 24-h light/dark period; each panel represents data from an individual monkey. Symbols are an average of 5 consecutive days including the 5 days immediately preceding discontinuation of Δ^9^-THC treatment (circles), the first and last 5 days of discontinued treatment (squares and diamonds, respectively), and the first 5 days of resumption of Δ^9^-THC treatment (triangles). The arrows indicate the time of Δ^9^-THC (1 mg/kg) or vehicle administration; vehicle was administered instead of Δ^9^-THC during the discontinuation phase. Below the abscissae the solid line denotes the dark period. Note the larger range of values on the ordinate of the lower panel as compared with the middle and top panels.

### Daily Δ^9^-THC treatment: discontinuation and resumption

On day 1 of discontinuation, there was a significant 2.6-fold increase in activity counts (261% of control; Figure 2, leftmost square). On day 2 activity counts were increased further, i.e., 2.8-fold, and remained significantly increased relative to control on days 3–6. Overall, there was a trend for activity to return to the pre-discontinuation control, i.e., 124% of control on day 20. Inspection of individual data in 1-h bins over the 24-h, light/dark period demonstrates that the increase in activity counts occurred predominantly during the light period (Figure 1, squares). The increase was evident at the beginning of the light period (7 AM) and throughout the day. When monkeys received Δ^9^-THC daily, they were relatively inactive by 7–8 pm (Figure 1, circles). During the first 5 days of discontinuation, however, activity remained high at 7–8 pm (Figure 1, squares). During the night, there was a trend for activity to be increased during brief intervals in two monkeys (Figure 1, squares over the black line in the middle and lowest panels) but not a third monkey (Figure 1 top).

**Figure 2.**
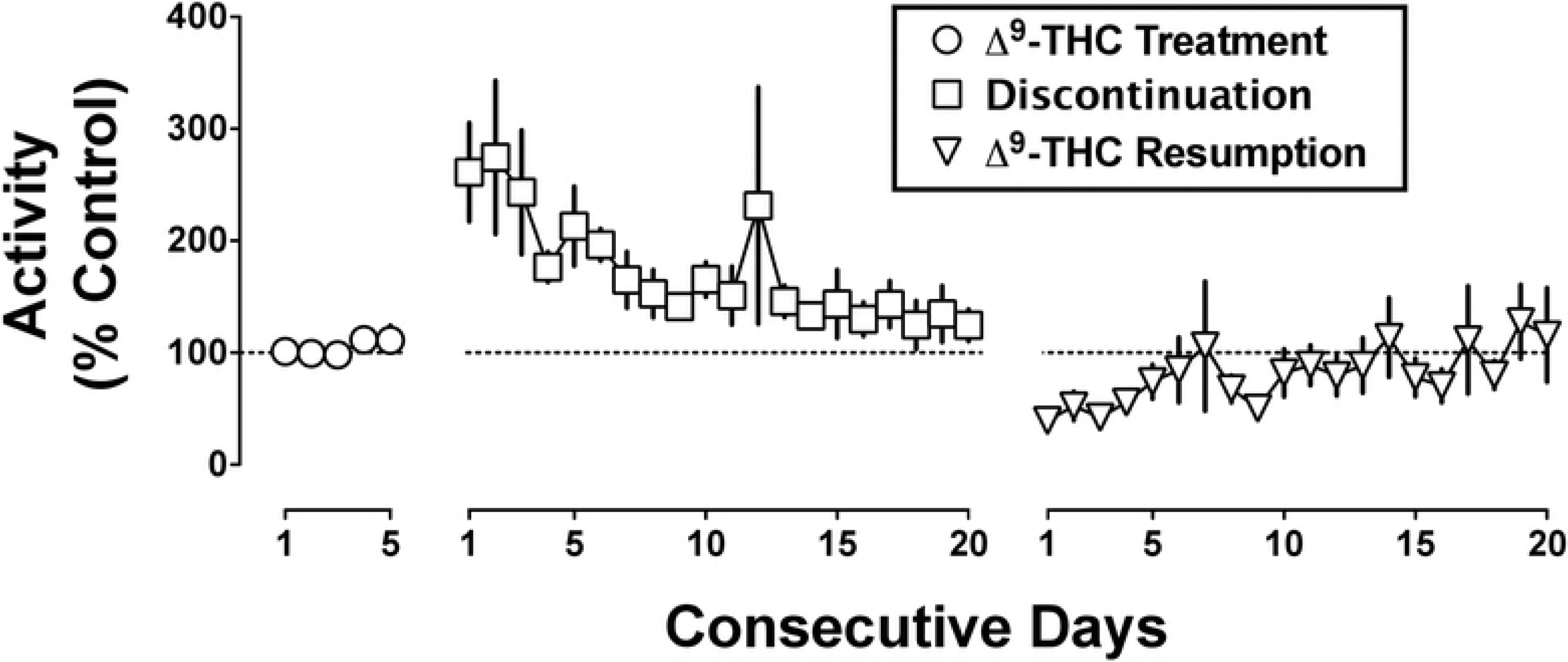
Activity counts cumulated daily (i.e., for 24 h) and expressed as a percentage of control as a function of consecutive days at various stages including before, during, and after discontinuation of Δ^9^-THC treatment. The control was defined as the mean activity counts of the 5 consecutive days immediately preceding discontinuation of Δ^9^-THC treatment. Symbols are the mean and error bars are the S.E.M.

Resumption of Δ^9^-THC treatment resulted in a marked decrease in activity counts (Figure 2, triangles). Activity was significantly decreased to 39% of control on day 1 and remained significantly decreased on days 2 and 3. Activity was no longer significantly different from control by day 4. The decreased activity upon resumption of Δ^9^-THC treatment was most prominent during the light period (Figure 1, triangles).

### Effects of Δ^9^-THC and rimonabant

In the five monkeys receiving chronic Δ^9^-THC (1 mg/kg/12 h), activity counts measured over a 2-h period following intravenous vehicle were 832, 3532, 3703, 6800, and 14433 for each respective monkey. In a separate group of monkeys (n=5) receiving intermittent Δ^9^-THC (0.1 mg/kg i.v. every 3 days on average), activity counts were somewhat higher, especially in two monkeys (4910, 5928, 15531, 25651, and 56430 for each respective monkey), as compared with monkeys receiving chronic Δ^9^-THC daily.

When expressed as a percentage of the vehicle control, intravenous rimonabant (0.1–3.2 mg/kg) dose-dependently increased activity in the chronic Δ^9^-THC group, but not the intermittent Δ^9^-THC group (F_2_,_36_=4.17; p<0.05). A dose of 1 mg/kg of rimonabant increased activity 13-fold in monkeys receiving chronic Δ^9^-THC (Figure 3 top left). The next larger dose (3.2 mg/kg) increased activity 20-fold; however, the variance was large due to activity being increased 70-fold in one monkey and unaltered in another monkey. The slopes of the Δ^9^-THC dose-response functions in the two groups did not significantly differ from each other. The intercepts also did not significantly differ; however, there was a tendency toward statistical significance (p=0.1) showing a difference in potency between the two groups and tolerance to Δ^9^-THC when its chronic treatment was resumed in the discontinuation study.

**Figure 3.**
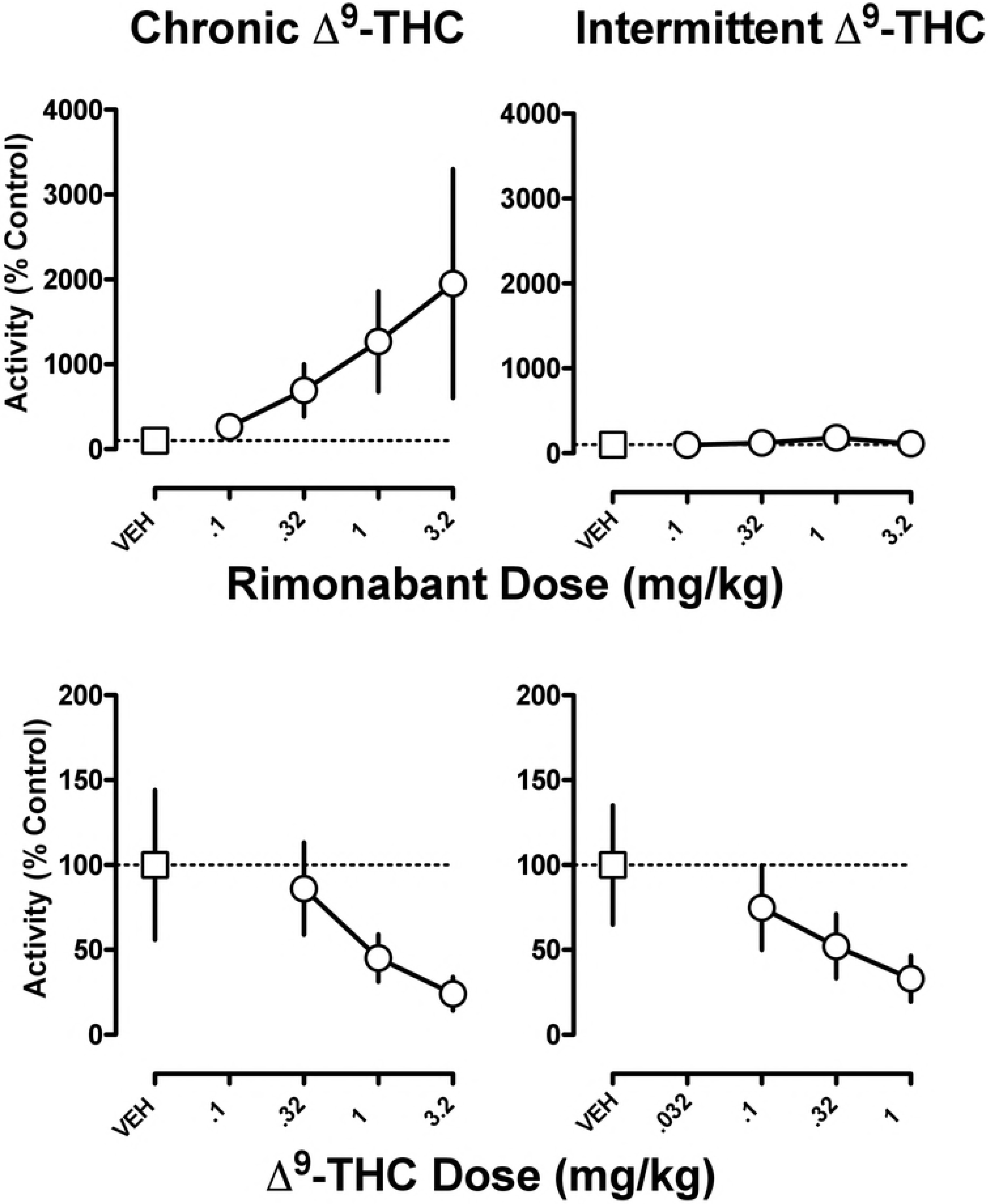
The effects of rimonabant (top panels) and Δ^9^-THC (bottom panels) in monkeys receiving either chronic Δ^9^-THC (left panels) or intermittent Δ^9^-THC (right panels). Activity counts are expressed as a percentage of the control calculated from the average of two separate vehicle tests. The abscissae denote dose in milligram per kilogram of body weight. Symbols are the mean and error bars are the S.E.M.

## DISCUSSION

Some advantages of the current study were the use of a standardized dosing regimen, a carefully controlled laboratory environment, systematic study of changes in behavior (home-cage activity) with a non-invasive device that continuously monitored activity, and use of non-human primates with similar drug and behavioral histories. Non-human primates are unique in that their biology is closely related to that of humans (Baskin 2013). However, expectation is not present in non-human animals and therefore cannot contribute to the changes in behavior and physiology comprising a drug withdrawal syndrome. In rhesus monkeys that had received 2 mg/kg of Δ^9^-THC daily for at least 2 years, abrupt discontinuation of treatment (i.e., replacing Δ^9^-Έ^ injections with vehicle injections) resulted in an immediate (i.e. within 24 h) increase in activity measured over a 24-h period that was apparent for several additional days. One interpretation is that, instead of withdrawal, the increased activity reflected a return to baseline from decreased activity induced by chronic Δ^9^-THC treatment. However, strong evidence to the contrary was demonstrated by the systematic decrease in hyperactivity back to the baseline measured during Δ^9^-THC treatment by day 20 of discontinuation. This time-limited increase in activity is consistent with a withdrawal syndrome, and the current time course is similar to that described for Δ^9^-THC withdrawal in humans (Hesse and Thylstrup 2013).

Looking at the pattern of activity expressed in 1-h bins across the 24-h period, increases in activity during discontinuation occurred primarily during the light portion, i.e. 0700–1700 h, of the light-dark cycle, a time when activity levels were already highest. While difficulty sleeping is a reliable component of cannabis withdrawal (Vandrey et al. 2011), any sleep disruption that might have occurred in monkeys was not evidenced by changes in activity during the night.

Administration of an antagonist can produce a withdrawal syndrome in agonist-dependent animals. The competitive cannabinoid CB_1_ receptor antagonist rimonabant has been used previously to produce Δ^9^-THC withdrawal in rodents and monkeys (Tsou et al. 1995; Stewart and McMahon 2010). Under the same conditions of Δ^9^-THC treatment used in the current study, a previous study reported a trend for increased activity after rimonabant (Stewart and McMahon 2010). In the current study, rimonabant produced a dose-dependent, marked increase in activity but only in monkeys receiving daily Δ^9^-THC treatment. The magnitude of increase was striking, i.e., 20-fold greater than the vehicle baseline. That the increased activity was not related to the direct effects of rimonabant was evidenced by the complete failure of rimonabant to increase activity in a separate group of monkeys that only received a relatively small dose of Δ^9^-THC (0.1 mg/kg i.v.) every 3 days on average. The marked increase in sensitivity to the effects of a cannabinoid antagonist in animals treated daily with Δ^9^-THC is consistent with a precipitated withdrawal syndrome. Sensitivity to the effects of rimonabant to decrease schedule-controlled responding was similarly increased in rhesus monkeys receiving the same chronic Δ^9^-THC treatment regimen (McMahon 2011), as well as in squirrel monkeys receiving chronic treatment with the cannabinoid agonist AM-411 (Desai et al. 2013). The qualitatively similar changes in activity observed not only after abrupt discontinuation of chronic Δ^9^-ΊΉ£, but also after rimonabant during Δ^9^-THC treatment, further suggest that withdrawal occurred under both conditions. However, the magnitude of change after rimonabant was markedly greater even though activity was cumulated over 2 h, as compared with 24 h in the discontinuation study. These data are consistent with the capacity of an antagonist to produce a greater magnitude of withdrawal as compared with abrupt discontinuation of treatment.

Δ^9^-THC typically decreases activity in rodents (Compton et al., 1996). Tolerance to the effects of Δ^9^-THC on activity in rhesus monkeys was evident when comparing the effects of daily Δ^9^-THC treatment before and after the discontinuation period. When Δ^9^-THC treatment was resumed after 20 days of discontinuation, activity was significantly decreased relative to the baseline calculated immediately prior to discontinuation of Δ^9^-THC treatment. However, within one week of resumed daily Δ^9^-THC treatment activity returned to levels in monkeys that had been receiving Δ^9^-THC treatment for years, indicative of tolerance. Intravenous Δ^9^-THC dose-dependently decreased activity in monkeys receiving chronic or intermittent Δ^9^-THC. While there was a tendency for Δ^9^-THC to be less potent in monkeys receiving chronic Δ^9^-THC versus intermittent Δ^9^-THC (Figure 3, compare circle above 0.32 mg/kg in bottom panels), the difference was not large enough to achieve statistical significance given the error variance and sample size. Nonetheless, the current results add to a substantial literature demonstrating tolerance to Δ^9^-THC (McMillan et al. 1970; Miczek 1979). The current results show that changes in activity in monkeys can be sensitive to both Δ^9^-THC tolerance and dependence, and are consistent with the observation that withdrawal signs are often opposite to the direct effects of the dependence-inducing drug.

Drug dependence can be operationally defined as a withdrawal syndrome that emerges upon abrupt discontinuation of chronic drug use. Many people are not aware of the dependence potential of cannabis or discount its importance, especially when compared with dependence and withdrawal resulting from chronic use of other abused drugs such as heroin or alcohol. Selfreported symptoms and observable signs of withdrawal that emerge in individuals who abruptly discontinue chronic cannabis use (e.g., smoking five cannabis cigarettes per day) include nervousness, irritability, depressed mood, sore joints, physical restlessness, and trouble sleeping (Haney 1999a; 1999b; Budney and Hughes 2006; Budney et al. 2008; Hesse and Thylstrup 2013). The clinical data have relied heavily on self-reports. While providing highly useful information on the characteristics of cannabinoid withdrawal, self-reports outside of the clinical laboratory cannot readily differentiate subtle changes in mood, perhaps influenced by expectation, that result from abrupt discontinuation of any habitual activity, from the physical dependence to cannabis that can be independently verified by systematic changes in behavior or physiology. Inpatient, double blind, placebo-controlled laboratory studies (e.g. Haney et al. 1999a; 1999b) can help to eliminate subject and experimenter bias.

In summary, these data provide strong evidence for physical dependence to Δ^9^-THC in non-human primates, as evidenced by withdrawal upon abrupt discontinuation of Δ^9^-THC treatment and administration of rimonabant during Δ^9^-THC treatment. The current results are consistent with the results of a previous study demonstrating a disruption of schedule-controlled responding in rhesus monkeys upon abrupt discontinuation of continuous intravenous infusion of Δ^9^-THC for at least 10 days (Beardsley et al., 1986). The disruption of schedule-controlled behavior also was time-limited and the time course was similar to that reported in the current study (i.e., 1–2 weeks in duration). Thus, Δ^9^-THC withdrawal is evidenced not only by a change in unlearned behavior (current study) but also learned behavior (Beardsley et al. 1986), suggesting that cannabinoid withdrawal can broadly disrupt behavior. However, withdrawal upon abrupt discontinuation of Δ^9^-THC is not always detected in rhesus monkeys when treatment duration is 3–9 weeks (Fredericks et al. 1981; McMahon, 2011), suggesting that Δ^9^-THC dependence is not always robust and might be more likely after lengthy periods of treatment. While the clinical significance of the withdrawal syndrome remains unclear, the current data demonstrate that physical dependence to Δ^9^-THC, and likely cannabis, clearly develops and must be recognized as a component of cannabis use disorders.

## Abbreviations

Δ^9^-THC: Δ^9^-tetrahydrocannabinol

## REFERENCES

Aceto MD, Scates SM, Lowe JA, Martin BR (1996) Dependence on delta 9-tetrahydrocannabinol: studies on precipitated and abrupt withdrawal. J Pharmacol Exp Ther 278:1290–1295

Baskin C (2013) The role and contributions of systems biology to the non-human primate model of influenza pathogenesis and vaccinology. Curr Top Microbiol Immunol 363:69–85

Beardsley PM, Balster RL, Harris LS (1986) Dependence on tetrahydrocannabinol in rhesus monkeys. J Pharmacol Exp Ther 239:311–319

Budney AJ, Hughes JR (2006) The cannabis withdrawal syndrome. Curr Opin Psychiatry 19:233–238

Budney AJ, Novy PL, Hughes JR (1999) Marijuana withdrawal among adults seeking treatment for marijuana dependence. Addiction 94:1311–1322

Budney AJ, Vandrey RG, Hughes JR, Thostenson JD, Bursac Z (2008) Comparison of cannabis and tobacco withdrawal: severity and contribution to relapse. J Subst Abuse Treat 35:362–368

Compton DR, Aceto MD, Lowe J, Martin BR (1996) In vivo characterization of a specific cannabinoid receptor antagonist (SR141716A): inhibition of delta 9-tetrahydrocannabinol-induced responses and apparent agonist activity. J Pharmacol Exp Ther 277:586–594

Desai RI, Thakur GA, Vemuri VK, Bajaj S, Makriyannis A, Bergman J (2013) Analysis of tolerance and behavioral/physical dependence during chronic CB1 agonist treatment: effects of CB1 agonists, antagonists, and noncannabinoid drugs. J Pharmacol Exp Ther 344:319–328

Fredericks AB, Benowitz NL (1980) An abstinence syndrome following chronic administration of delta-9-terahydrocannabinol in rhesus monkeys. Psychopharmacology 71:201–202

Fredericks AB, Benowitz NL, Savanapridi CY (1981) The cardiovascular and autonomic effects of repeated administration of delta-9-tetrahydrocannabinol to rhesus monkeys. J Pharmacol Exp Ther 216:247–253

Haney M, Ward AS, Comer SD, Foltin RW, Fischman MW (1999a) Abstinence symptoms following smoked marijuana in humans. Psychopharmacology 141:395–404

Haney M, Ward AS, Comer SD, Foltin RW, Fischman MW (1999b) Abstinence symptoms following oral THC administration to humans. Psychopharmacology 141:385–394

Harris RT, Waters W, McLendon D (1974) Evaluation of reinforcing capability of delta-9-tetrahydrocannabinol in rhesus monkeys. Psychopharmacologia 37:23–29

Hesse M, Thylstrup B (2013) Time course of DSM-5 cannabis withdrawal symptoms in poly-substance abusers. BMC Psychiatry 13:258

Institute for Laboratory Animal Research (2011) Guide for the care and use of laboratory animals, 8th edn. Institute for Laboratory Animal Research, Division of Earth and Life Sciences, National Research Council, Washington, DC

Jones RT (1983) Cannabis and health. Annu Rev Med 34:247–258

McMahon LR (2011) Chronic Δ^9^-tetrahydrocannabinol treatment in rhesus monkeys: differential tolerance and cross-tolerance among cannabinoids. Br J Pharmacol 162:1060–1073

McMillan DE, Harris LS, Frankenheim JM, Kennedy JS (1970) l-D9-trans-tetrahydrocannabinol in pigeons: tolerance to the behavioral effects. Science 169:501–503

Miczek KA (1979) Chronic D9-tetrahydrocannabinol in rats: effect on social interactions, mouse killing, motor activity, consummatory behavior, and body temperature. Psychopharmacology 60:137–146

Stewart JL, McMahon LR (2010) Rimonabant-induced Delta9-tetrahydrocannabinol withdrawal in rhesus monkeys: discriminative stimulus effects and other withdrawal signs. J Pharmacol Exp Ther 334:347–356

Tsou K, Patrick SL, Walker JM (1995) Physical withdrawal in rats tolerant to delta 9-tetrahydrocannabinol precipitated by a cannabinoid receptor antagonist. Eur J Pharmacol 280:R13–R15

Vandrey RG, Budney AJ, Hughes JR, Liguori A (2008) A within-subject comparison of withdrawal symptoms during abstinence from cannabis, tobacco, and both substances. Drug Alcohol Depend 92:48–54

Vandrey R, Haney M (2009) Pharmacotherapy for cannabis dependence: how close are we? CNS Drugs 23:543–553

Vandrey R, Smith MT, McCann UD, Budney AJ, Curran EM (2011) Sleep disturbance and the effects of extended-release zolpidem during cannabis withdrawal. Drug Alcohol Depend 117:38–44

Young AM, Katz JL, Woods JH (1981) Behavioral effects of levonantradol and nantradol in the rhesus monkey. J Clin Pharmacol 21:348S–360S

